# Indirect actuation reduces flight power requirements in *Manduca sexta* via elastic energy exchange

**DOI:** 10.1101/727743

**Authors:** Jeff Gau, Nick Gravish, Simon Sponberg

## Abstract

In the vast majority of flying insects, wing movements are generated indirectly via the deformations of the exoskeleton. Indirect measurements of inertial and aerodynamic power requirements suggest that elastic energy exchange in spring-like structures may reduce the high power requirements of flight by recovering energy from one wingstroke to the next. We directly measured deformation mechanics and elastic energy storage in a hawkmoth *Manduca sexta* thorax by recording the force required to deform the thorax over a frequency range encompassing typical wingbeat frequencies. We found that a structural damping model, not a viscoelastic model, accurately describes the thorax’s linear spring-like properties and frequency independent dissipation. The energy recovered from thorax deformations is sufficient to minimize flight power requirements. By removing the passive musculature, we find that the exoskeleton determines thorax mechanics. To assess the factors that determine the exoskeleton’s spring-like properties, we isolated functional thorax regions, disrupted strain in an otherwise intact thorax, and compared results to a homogeneous hemisphere. We found that mechanical coupling between spatially separated thorax regions improves energy exchange performance. Furthermore, local mechanical properties depend on global strain patterns. Finally, the addition of scutum deformations via indirect actuation provides additional energy recovery without added dissipation.

## 1 Introduction

Indirect actuation is a prevalent feature among flying insects. Unlike traditional skeletal muscle, the power muscles in many species indirectly generate wing movements by deforming a continuous exoskeleton surrounding the thorax [1]. Like the complex interactions between power limited muscles, non-ideal latches, and imperfect springs in power amplified biological systems, the introduction of the deformable exoskeleton could have significant consequences on flight mechanics [2]. For instance, the thorax’s frequency response may cause some wingbeat frequencies to be energetically favorable, thereby encouraging flight within a narrow band as seen in *Manduca sexta* [3]. In addition, for the past 60 years, it has been thought that these exoskeletal deformations may reduce the energy required for flight by providing elastic energy storage and return. However, it is unclear what factors give rise to spring-like properties in a materially heterogeneous structure with complex geometry. Despite the prevalence of indirect actuation in insect flight, it is unknown how the integrated mechanical system responds to changes in wingbeat frequencies, how thorax mechanics affect flight energetics, and what factors give rise to the thorax’s spring-like properties.

To fly, insects must provide power to accelerate and decelerate their oscillating wings (*P_inertial_*) as well as to move the surrounding air (*P_aero_*). Assuming symmetric upstroke and downstrokes, no energetic costs of wing deceleration, *P_aero_* < *P_inertial_*, and zero elastic energy exchange, the total power required for flight is 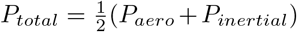 [4, 5]. Under these conditions, the insect must only provide mechanical power during the acceleration phase of each half stroke because *P_inertial_* is converted into *P_aero_* during deceleration. If there is elastic energy exchange, excess wing kinetic energy can be absorbed and returned by elastic elements in the flight system (*P_return_*). In an ideal case, *P_total_* = *P_aero_*. Because *P_aero_* < *P_inertial_*, this corresponds to a net reduction in *P_total_*. In this case, all excess inertial power (*P_inertial_* – *P_aero_*) is stored in spring-like structures and returned to reduce *P_inertial_* for the subsequent half stroke [4, 5].

Several modeling and indirect mechanical measures have estimated the potential benefits of elastic energy exchange afforded by the thorax and muscles during flight [6, 7, 8, 9]. Blade element models, computational fluid dynamics, and tomographic particle image velocimetry estimates suggest that perfect elastic energy exchange could reduce *P_total_* by up to 20 - 35% in *Manduca sexta* [8, 9, 7]. These estimates indicate the potential benefits of elastic energy storage in flapping wing systems. However, there is a lack of direct mechanical measurements of the insect thorax to determine if elastic energy storage and recovery occurs during flight.

There are several potential sources of elastic energy exchange in insect flight. The antagonistic extension of elastic elements in both passive and active muscle as well as exoskeletal deformations may store and return elastic energy (Fig. 1 b) [10, 11, 12, 13]. In particular, temperature gradients in *Manduca sexta* enables crossbridges to remain bound and function as springs, although the energy exchange capacity has not been quantified [13]. Unlike muscle, it is unclear what factors would enable substantial elastic energy exchange in the exoskeleton. While the exoskeleton is composed of resilient materials, such as resilin and chitin, material properties are not the sole determinant of bulk mechanical properties [14, 15, 16, 17]. For instance, mechanical coupling of bending and stretching due to a structure’s shape can significantly alter bulk mechanics [18]. In addition, large amplitude heterogeneous strain may concentrate deformations in regions with unfavorable properties for energy exchange (Fig. 2 d) [19]. Thus, interactions between exoskeletal shape and material composition are significant for determining elastic energy exchange capacity. While all three sources may recycle elastic energy, there is a maximum useful elastic energy return beyond which *P_total_* is not affected.

**Figure 1:**
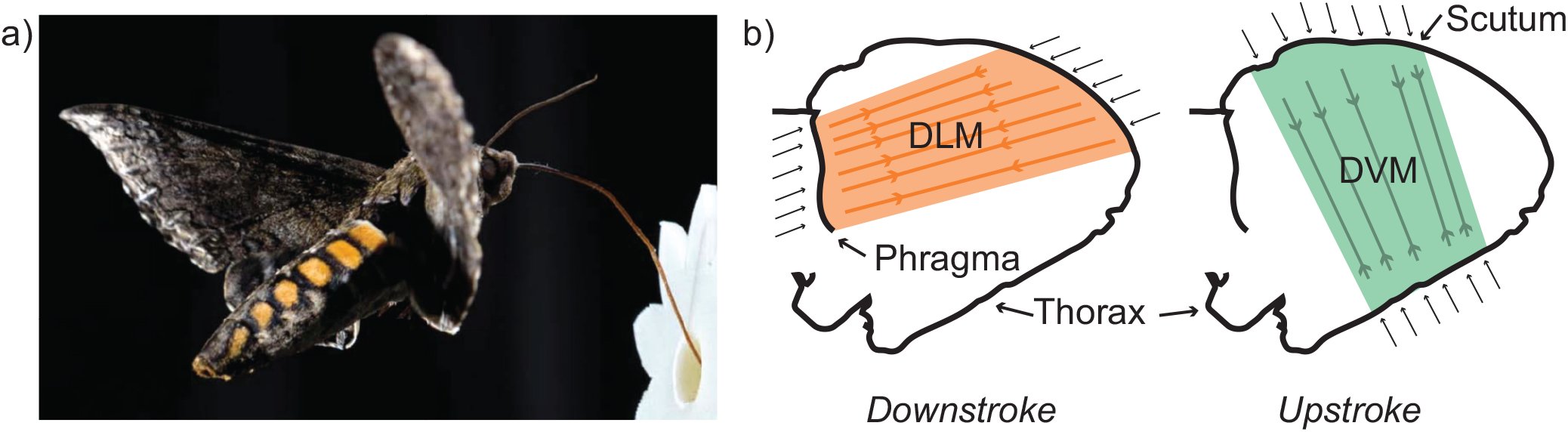
Insects with indirect actuation deform their exoskeleton to move their wings. a) The hawkmoth *Manduca sexta* utilizes indirect actuation to fly. b) Two sets of muscles, the dorsolongitudinal (DLM) and dorsoventral (DVM) attach to the thorax. The DLM attaches to the posterior phragma, which is an insertion of the exoskeleton into the thorax. The DVM attaches to the scutum, a smooth plate on the top of the thorax. The DLM and DVM act against each other to drive the upstroke and downstroke movements by deforming the thorax.

**Figure 2:**
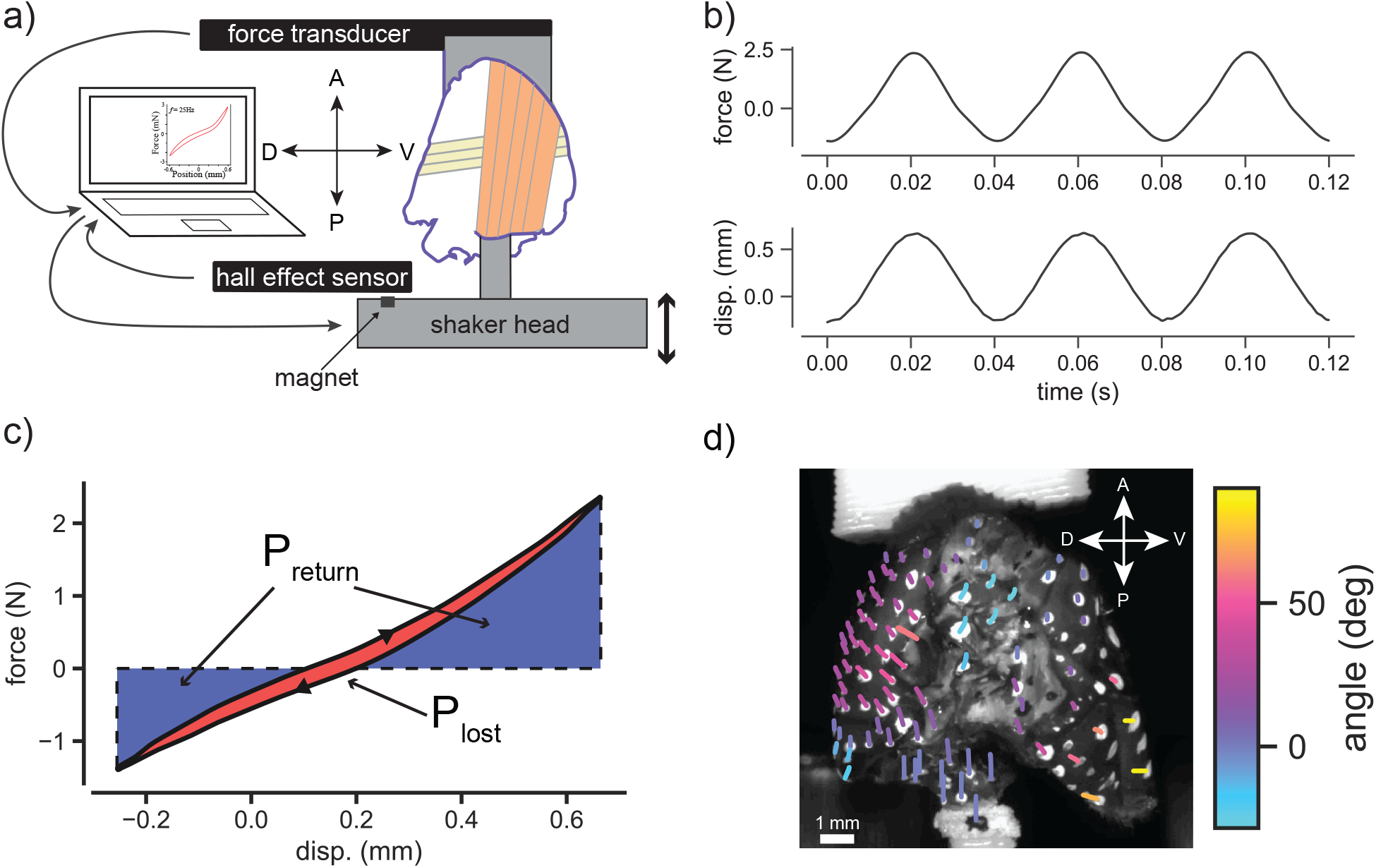
Experimental overview. a) Experimental setup. The thorax is mounted between a dynamic mechanical testing device and a force transducer. We prescribe a sinusoidal length input to physiological amplitudes and measure the force required to deform the thorax. b) Representative force and displacement measurements from an intact thorax with passive musculature at 25 Hz. c) Force-displacement plot from the same data as b), but averaged over all oscillations. Blue area denotes *P_return_* while red area denotes *P_lost_*. The displacements are asymmetric because the operating length of downstroke muscles is 0.21 mm shorter than rest length [22]. d) Visualization of strain through the second thoracic segment during a physiological oscillation at 1 Hz. We recorded high speed video at 200 fps. White and black dots were painted on the thorax to aid in tracking. Color denotes deformation angle off of the vertical while traces highlight the movement of each tracked point.

In addition to energy recovery, energy loss during thorax deformations may substantially alter flight power requirements. Under the large strain conditions (9% peak-to-peak) observed in *Manduca sexta*, biological springs are often far from ideal [20]. Instead, they typically introduce substantial, and often, frequency dependent damping [17, 21]. Similar to *P_aero_*, any dissipation in the thorax is unrecoverable. If *P_lost_* exceeds *P_return_*, then the net effect of thorax deformations is an increase in *P_total_*. Furthermore, if damping is frequency dependent as seen in passive and active flight muscle [21], then there may be energetically preferable wingbeat frequencies and possible limits on the range of energetically attainable wingbeat frequencies.

We hypothesize that the thorax acts as a damped spring that substantially reduces *P_total_*. To test this hypothesis, we employ a custom vibration apparatus to measure the thorax’s passive mechanical properties under physiological conditions. We provide an oscillatory displacement to the thorax, mimicking the action of the flight power muscles. We measured the force required to deform the thorax to determine elastic energy storage and recovery in a typical wingbeat. Through frequency sweeps encompassing the typical wing beat frequency, we develop a model of thorax elasticity and damping. In addition, we employ a series of thorax manipulations to identify how different exoskeleton regions contribute to flight energetics. We compare these manipulations to an ideal homogeneous hemisphere (ping pong ball) to assess the effects of shape and material. Lastly, we repeat mechanical testing experiments on a thorax with removed musculature to determine the relative importance of passive muscles on flight energetics.

## 2 Methods

### 2.1 Animals

*Manduca sexta* used in these experiments were obtained as pupae from colonies maintained at the University of Washington and Case Western Reserve University. Moths were kept on a 12h:12h day:night cycle. We used 7 males and 27 females between 1 and 7 days post-eclosion. Moth mass equaled 2.13 ± 0.47 g.

### 2.2 Thorax preparation

Following a 30 minute cold anesthesia, we removed the head, abdomen, wings, and legs from each moth to isolate the thorax. In some preparations (N = 10), we removed the flight musculature, while in others we left the muscle intact (N = 24). For the intact muscle preparations, we silenced neural activity by cutting the outward projections of the thoracic ganglion to the flight musculature. We then removed the first thoracic section, leaving only the thorax segments involved in flight (pterothorax). Finally, we used damp wipes and compressed air to remove scales from the exoskeleton.

#### 2.2.1 Muscle Removal

The anterior and posterior ends of the pterothorax have large openings to remove muscle. We used forceps and scalpels to remove flight muscle from the pterothorax. We ensured that we removed at least 0.1 g of muscle, which is in rough agreement with previous reports of DLM muscle mass [20]. Any remaining pieces of muscle were negligible because we ensured that they only had one intact attachment point. Therefore, these muscle remnants could not be stretched and generate significant force during our experiments.

### 2.3 Thorax manipulations

#### 2.3.1 Strain disruption cuts

For these manipulations, we used a series of cuts to disrupt strain through the exoskeleton without removing material (Fig. 7 a & b). The objective of these cuts was to disrupt the transverse arch, which is known to increase longitudinal stiffness. For the longitudinal cut condition, we used a razor blade to cut a single anterior-posterior cut along the midline of the scutum. We then performed the triple longitudinal cut condition in which two parallel cuts are made on either side of the first longitudinal cut. This created three anterior-posterior cuts on the scutum.

#### 2.3.2 Wing joint and scutum isolation cuts

These cuts removed material in the exoskeleton to isolate strain to certain functional regions (Fig. 6 a & b). For the isolated joint condition, we removed the scutum to isolate strain to the wing joint. For the isolated scutum condition, we removed the wing joint to isolate strain to the scutum. The total material displaced in the isolated joint plus isolated scutum conditions should equal that of an intact thorax because the dorsal regions of the thorax are largely decoupled from the ventral.

### 2.4 Shaker preparation

We used cyanoacrylate glue to secure a 3D printed shaft to the posterior phragma (Fig. 1 b and Fig. 2 A & D). This shaft was then mounted to the head of a custom shaker mechanism as described in [23]. On the anterior end, we used cyanoacrylate to rigidly attach the anterior phragma to a cantilever beam force transducer rated to 10 N with a resonance frequency of 300 Hz (FORT1000, World Precision Instruments, Sarasota, FL). The force transducer itself was mounted to a 3-axis micromanipulator, which enabled us to precisely set the rest length of the thorax. Using the force transducer, we weighed the ABS attachment pieces plus cyanoacralyte. After mounting the thorax vertically, we returned the force to match this weight. The micromanipulator also allowed us to precompress the thorax to physiological amplitudes [22]. To measure thorax compression and tension, we used an analog hall effect sensor (DVR5053-Q1, Texas Instruments, Dallas, Tx). This sensor was calibrated to measure the position of a permanent magnet attached to the shaker head. We drove the shaker head with an electrodynamic vibration testing system (VTS 600, Vibration Test Systems, Aurora, OH). Critically, the anterior and posterior phragmas are the structures to which the downstroke power muscles attach *in vivo*. Attaching our experimental apparatus to these structures and displacing along the contraction axis of the main downstroke muscle (DLM) axis ensured physiologically relevant deformations.

### 2.5 Dynamic mechanical testing

After procompressing the thorax, we prescribed sinusoidal displacements to a physiological amplitude of 0.46 mm over a frequency sweep from 0.1 to 90 Hz [22]. This frequency range encompasses the hawkmoth’s wingbeat frequency of 25 Hz. At each frequency, we ensured the thorax reached steady state and simultaneously measured forces and displacements, as described above.

### 2.6 Strain mapping

In one thorax, we mapped 2D strain in the parasagittal plane. We removed the third thoracic segment to visualize the phragma and left the flight muscle intact. We then painted small white and black dots on the thorax to aid strain visualization (Fig. 2 d). We deformed the thorax at 1 Hz while recording from from a machine vision camera at 200 Hz (BlackFly S BFS-U3-4S2M-CS, FLIR Integrated Imaging Solutions, Inc. Richmond, BC, Canada).

### 2.7 Ping pong ball preparation

We repeated the analogous procedures to prepare ping pong balls. We chose ping pong balls because they are highly uniform, precisely spherical, and made of a material, celluloid, known to have structurally damped properties [24]. Because ping pong balls are spherical and homogeneous, there is no true “scutum” or “wing hinge”. We therefore visually determined analogous “scutum” and “wing hinge regions. Similar to the thorax, we used ABS shafts to attach the ping pong ball to the custom shaker. For dynamic material testing, we used a displacement of 0.5 mm and precompression of 0.2 mm. We found that ping pong balls had similar peak forces to the thoraxes, indicating the the thorax and ping pong ball experiments were in similar dynamic regimes for the shaker apparatus.

### 2.8 Desiccation verification

To assess the effects of desiccation on thorax properties, we report *P_return_* and *P_lost_* on three thoraxes with passive musculature (Fig. S1). Following the dynamic mechanical testing procedure outlined above, we repeatedly measured *P_return_* and *P_lost_* to assess the effects of desiccation on elastic energy exchange. We found that both *P_return_* and *P_lost_* increased with time, but that the conclusions in this report hold true regardless of desiccation effects.

### 2.9 Data analysis

#### 2.9.1 Empirical calculations of power exchange

We calculated three body mass-specific powers: power required to drive the displacements 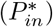, power returned via elastic energy storage and return 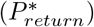, and power dissipated during an oscillation 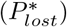. Because the wings have been removed, 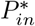 is just the power necessary to deform the thorax and accelerate thoracic mass. 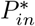 is the sum of 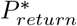 and 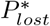. We begin by empirically integrating force over displacement to determine the analogous energy metrics (Fig. 2 c). We multiply energy by oscillation frequency to arrive at power. We then normalize by body mass to arrive at body mass-specific powers (*P*\*). Finally, we calculate resilience (R), which is a measure of spring efficiency, as

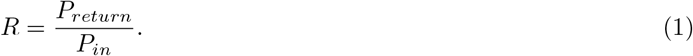

#### 2.9.2 Thorax elastic linearity

To assess linearity, we compare resilience values between those derived from empirical force-displacement data versus resilience values of a linear fit to the force-displacement data (Fig. 4 D). For the linear fit, force and displacement amplitude are calculated as the maximum amplitude of the corresponding Fourier transforms. Phase lag between force and displacement (*ϕ*) is determined by 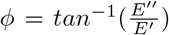. From the phase lag, *E*, and strain measurements, we generate a force-displacement ellipse. Because precompression affects *R*, we offset this ellipse by the force and displacement offsets measured in the thoraxes. Finally, we empirically integrate to determine *R*.

#### 2.9.3 Statistics

To determine the frequency dependence of data, we used a linear mixed effects model [25]. In this model, each trial was represented as a random variable. To compare data at wingbeat frequency, we used two-sided t-tests. All reported errors are one standard deviation.

## 3 Results

### 3.1 The thorax behaves like a linear, structurally damped spring

The force-displacement traces provide a preliminary assessment of the thorax’s mechanical properties (Fig. 3 a & b). The thorax is stiff, requiring 2 N to compress under physiological conditions. This force magnitude agrees with muscle force output previously reported for *Manduca sexta* [20]. Despite its structural complexity, the thorax is well-approximated as a linear material, both with and without passive musculature. Under both conditions, there is evidence of *P_return_* and *P_lost_* in the thorax. Because we precompressed the thoraxes to physiological conditions, the majority of the elastic energy returned occurs during the down stroke. Finally, for both the passive and removed musculature conditions, the force-displacement relationship is largely frequency independent, with only visible differences evident between the 0.1 and 90 Hz trials. From mixed linear model regression, we found that force does increases very shallowly by 1.693 ± 0.11 mN per Hz (*p* < 0.001) and resilience decreases by −0.052 ± 0.002 % per Hz (*p* < 0.001) in thoraxes with passive musculature. This indicates that stiffness and damping are weakly frequency dependent over this range, but that the magnitude of change from 0.1 to 90 Hz (9% for stiffness and 6% for resilience) is likely biologically insignificant, especially because *Manduca sexta* fly within a narrow frequency range around 25 Hz [3].

**Figure 3:**
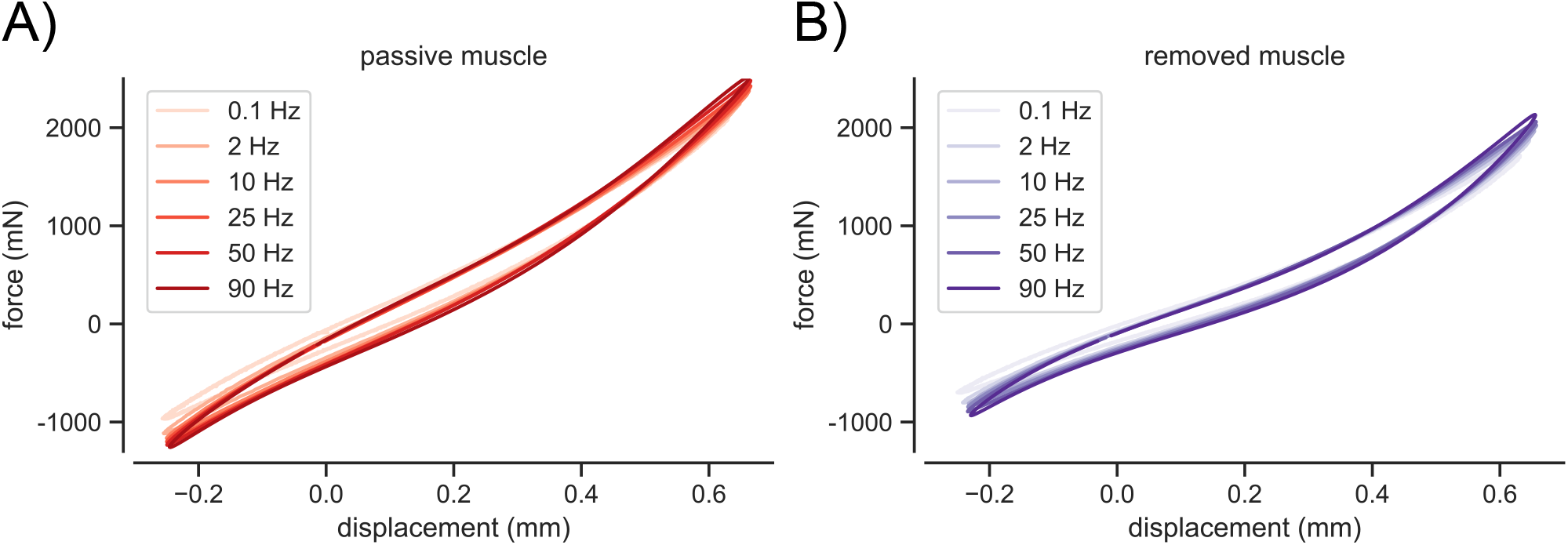
Force-displacement traces at representative frequencies from 0.1 to 90 Hz for an intact thorax with a) passive muscle (N = 24) and b) removed muscle (N = 10). For each frequency, the traces represent the average force and average displacement at each time point. Positive displacement denotes compression.

To better characterize these observations, we sought a mathematical model of thorax elasticity and damping. A linear force-displacement model requires a minimum of two parameters (storage and loss). Therefore, we considered two classical two-parameter models. The Kelvin-Voigt model consists of a parallel spring and viscous damper (Fig. 4 a, inset), as defined in Eq. 2, where *m* is the mass of the object, *c* is the damping coefficient, and *k* is the spring stiffness. The damping force in this system is velocity dependent.

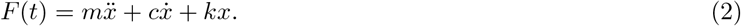

**Figure 4:**
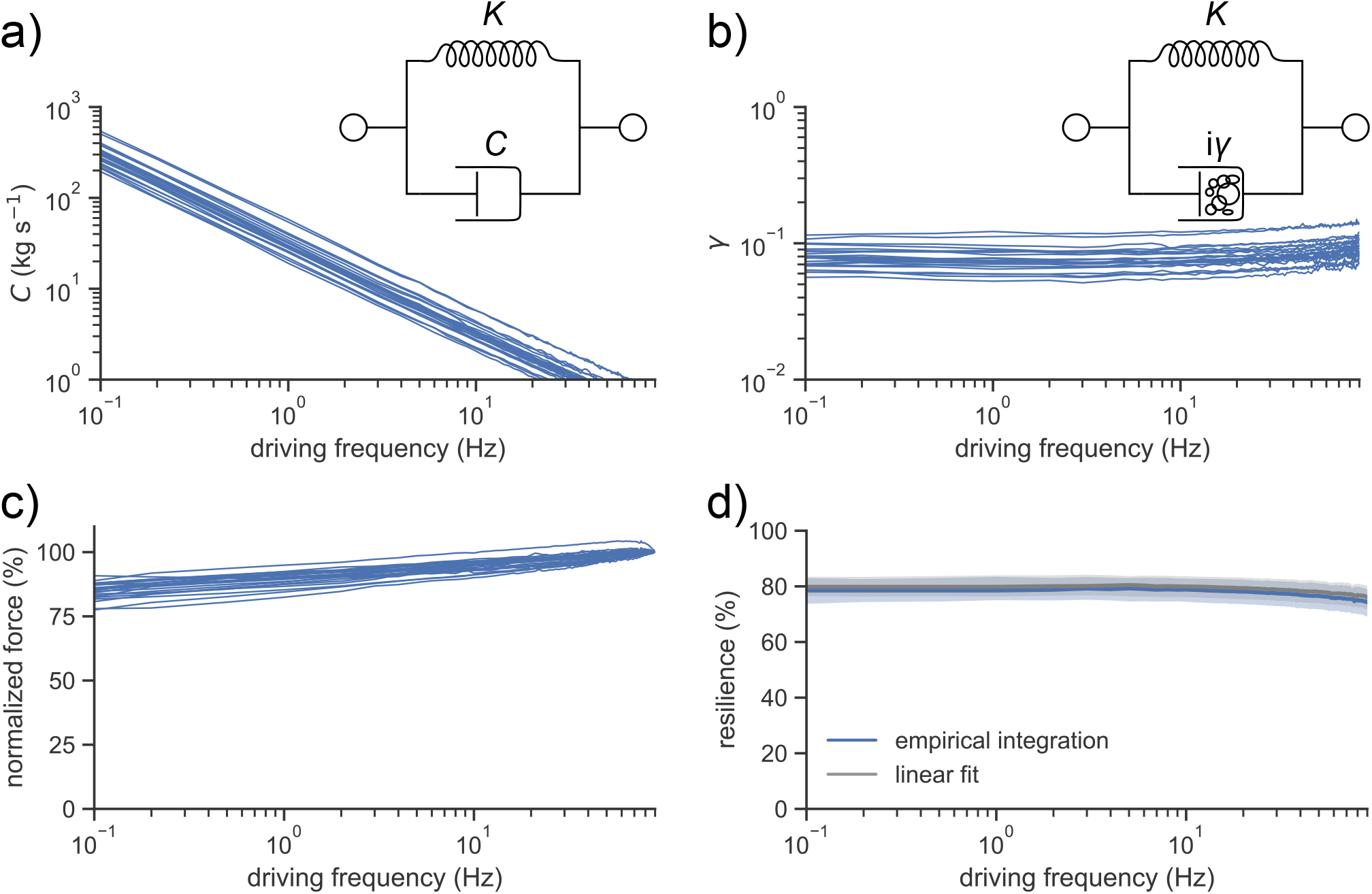
Frequency response and mechanical characterization of intact thorax with passive musculature (N = 24). Each line denotes one individual. a) Damping coefficient *c* for a Kelvin-Voigt model fit to experimental data. b) Damping coefficient *γ* for a structural damping model fit to experimental data. c) Normalized peak force required to deform the thorax versus frequency. For each individual, force is normalized to the peak force at 90 Hz. d) Resilience versus calculated for experimental force-displacement data (blue) and for a linear fit (gray). Shaded area denotes one standard deviation above and below the mean.

Next, we considered a structural damping model (Fig. 4 b, inset and Eq. 3) because it accurately characterized the bending of cockroach legs over a wide frequency range [26]. In this model, *m* is the mass of the object, *γ* is the structural damping coefficient, and *k* is the spring stiffness. In this model, damping force is position dependent.

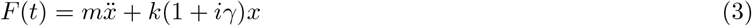

Both models are linear viscoelastic models but have significantly different frequency predictions. To determine the efficacy of each model, we assumed negligible mass and fit the stiffness *k* and loss coefficient (*c* or *γ*) at each oscillation frequency.

To calculate the structural damping coefficient (*γ*), the Kelvin-Voigt damping coefficient *c*, and Young’s Modulus *E*, we first determine the storage modulus E’ and the loss modulus E”. We took the Fourier transform of stress and strain at each oscillation frequency. We then divided the peak complex stress by peak complex strain to determine the complex modulus, where E’ is the real portion and E” is the imaginary component. Assuming *m* is negligible, *E* = *E*′. Next we calculated the damping coefficients as 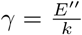 and 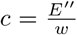, where w is the angular frequency of oscillation [26].

The Kelvin-Voigt damping parameter, *c* changed by nearly three-orders of magnitude over the driving frequency range (Fig. 4 a). The strong frequency dependence of *c* indicates that the Kelvin-Voigt model is a poor representation of thorax mechanics because the thorax itself is not changing as a function of oscillation frequency. In contrast, the structural damping parameter *γ* was nearly constant over the entire driving frequency range (Fig. 4 c). These results suggest that the thorax is best described by a structural damping model because this model has frequency independent parameters.

There is strong agreement between resilience values from empirical force-displacement data and resilience values calculated from a linear fit of the same data (Fig. 4 d). This suggest that deviations from linearity observed in the force-displacement data (Fig. 4 a) do not significantly affect energetics. In summary, we conclude that the thorax behaves as a structurally damped, linear spring.

### 3.2 Passive muscle is negligible for elastic energy exchange

To assess the relative contributions of the exoskeleton and passive muscle on thorax elasticity under physiological conditions, we compared power measurements for thoraxes with passive (N = 24) and removed (N = 10) musculature. We found that removing the passive musculature significantly reduced the peak force to deform the thorax from 1730 ± 430 mN to 1350 ± 250 mN (*p* = 0.015) (Fig. 5 a). However, there was no statistical difference in resilience values between thoraxes with passive and removed musculature (*p* = 0.277) (Fig. 5 b). Although passive muscle added significant stiffness, when we divided the force-displacement plots into the respective power metrics and normalized by body mass, we found no statistically significant difference in 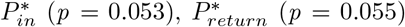, or 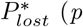 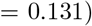 (Fig. 5 c). These results suggest that the passive thorax’s energy exchange capacity is dominated by the exoskeleton and passive muscle is negligible. However, activating muscle can engage additional elastic elements, such as crossbridges, to increase thorax stiffness [13].

**Figure 5:**
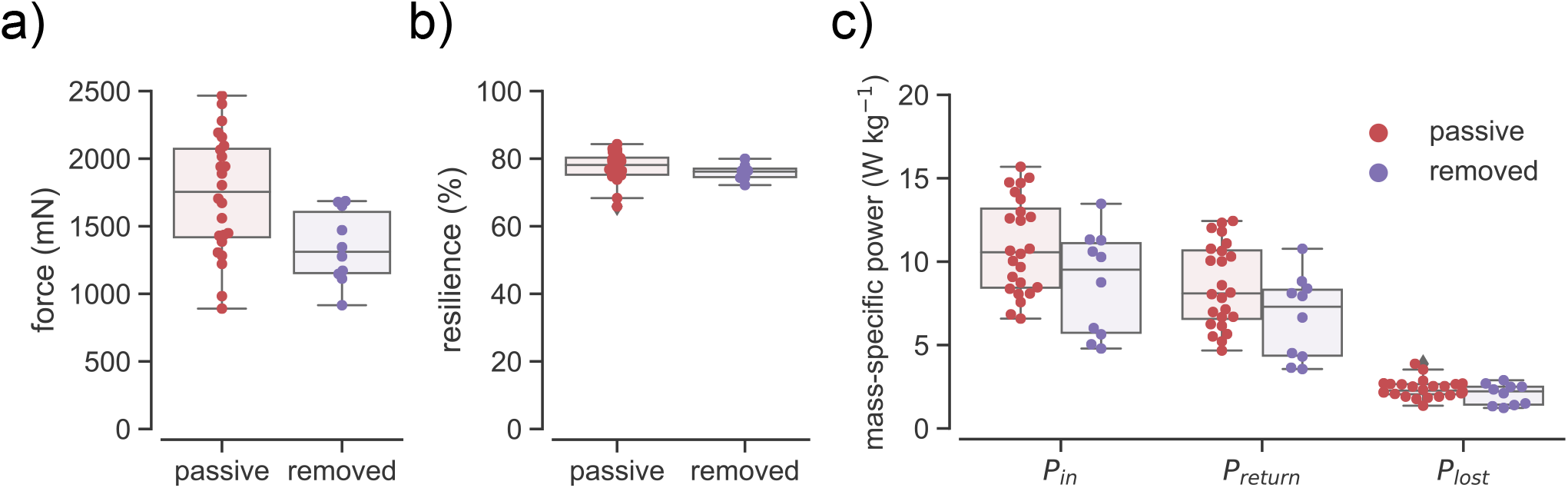
Mechanics of passive muscle are negligible. All data was taken at oscillation frequencies at 25 Hz. Boxplots denote the mean, quartiles, and range of the raw data while circles are individual data points. a) Peak force required to deform the thorax with passive (N = 24) and removed (N = 10) musculature. b) Resilience for thoraxes with passive and removed musculature. c) Mass-specific power measures for thoraxes with passive and removed musculature.

Furthermore, the high stiffness and resilience of the passive thorax indicates substantial elastic energy exchange under physiological conditions. With passive musculature, we obtained a *P_in_* of 10.9 ± 2.8 W kg^−1^, *P_return_* of 8.5 ± 2.4 W kg^−1^, and a *P_lost_* of 2.4 ± 0.5 W kg^−1^. With muscle removed, we found a *P_in_* of 8.7 ± 3.0, *P_return_* of 6.7 ± 2.4, and *P_lost_* of 2.1 ± 0.6 W kg^−1^.

### 3.3 Isolation of thorax regions reduces regional energy exchange performance

From visualizations of thorax strain patterns (Fig. 2 d), it is evident that the wing joint and scutum move orthogonally as the thorax is stretched and compressed (Fig. 6 a & b). To assess how interactions between these regions affect bulk mechanics, we performed dynamic mechanical testing on thoraxes with isolated wing joints and isolated scutums. We performed analogous experiments on a homogeneous hemisphere to ground our interpretation of these results. We used ping pong balls because they are homogeneous, structurally damped hemispheres [24]. Although ping pong balls do not naturally possess scutums or wing hinges, we removed the analogous regions to gain intuition on how a thorax with reduced complexity might behave.

**Figure 6:**
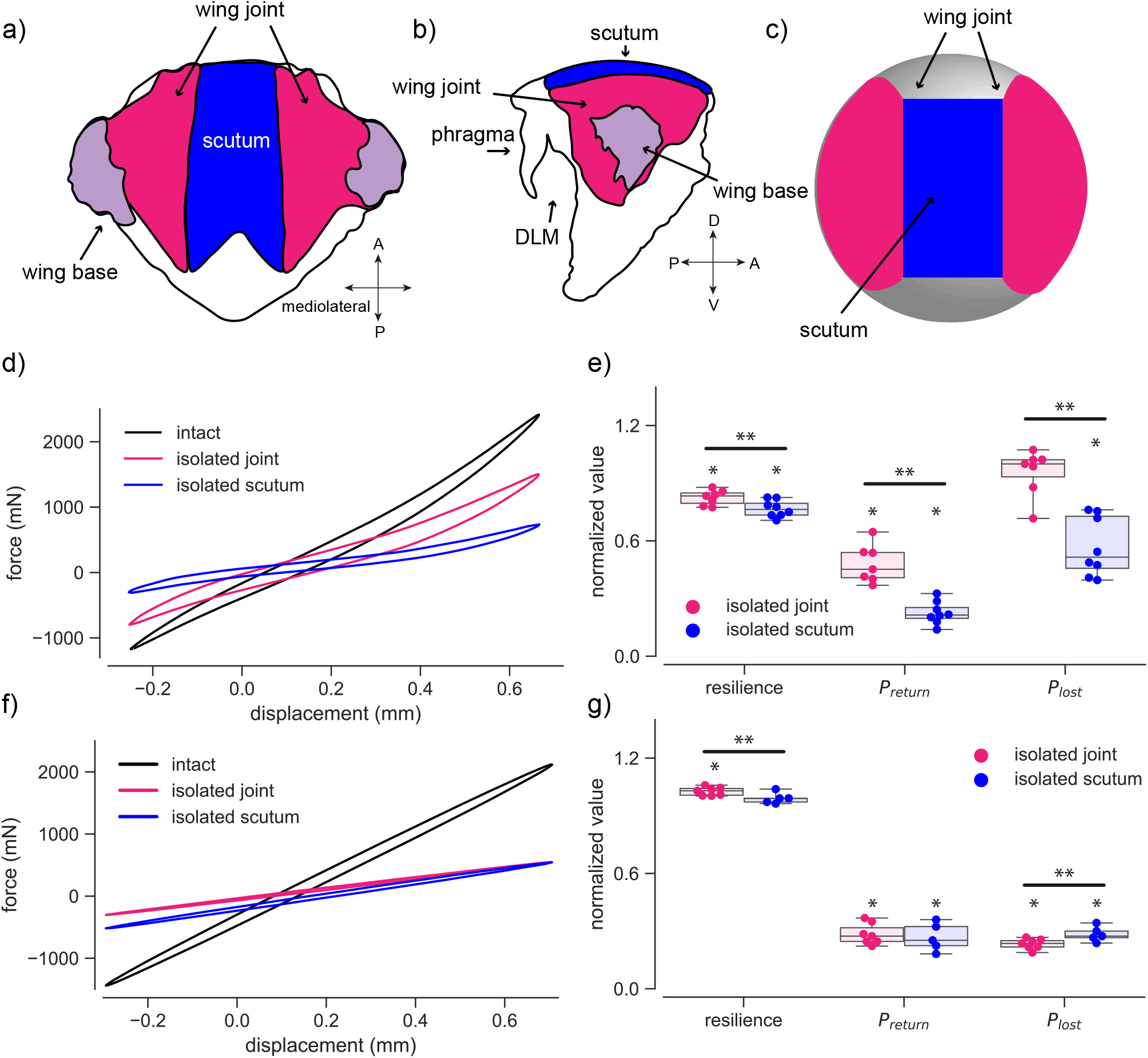
Region isolation experiments. a) Dorsal view of the second thoracic segment, which contains the majority of flight muscle and forewings. The wing joint is in pink, scutum in blue, and wing base in purple. b) Lateral view of the second thoracic segment. In addition to the regions labeled in a), the phragma is visible. c) Analogous dorsal view of a ping pong ball highlighting the regions labeled in a). d) Representative force-displacement plots for an intact, isolated joint, and isolated scutum thorax. At each time point, we plot the average force against the average displacement for each condition. Positive displacement denotes compression. e) Effects of regions isolation on resilience, *P_return_*, and *P_lost_* in thoraxes with passive musculature and isolated scutums (N = 8) or isolated wing joints (N = 7). All data was normalized to the paired, intact thorax and taken at an oscillation frequency of 25 Hz. Boxplots denote the mean, quartiles, and range of the raw data while circles are individual data points. Single * denote a significant difference from the paired, intact thorax while ** mark differences between conditions. f) Same as d) but for a ping pong ball. g) Same measures as e) but for ping pong balls with isolated scutums (N = 7) or isolated joints (N = 5).

**Figure 7:**
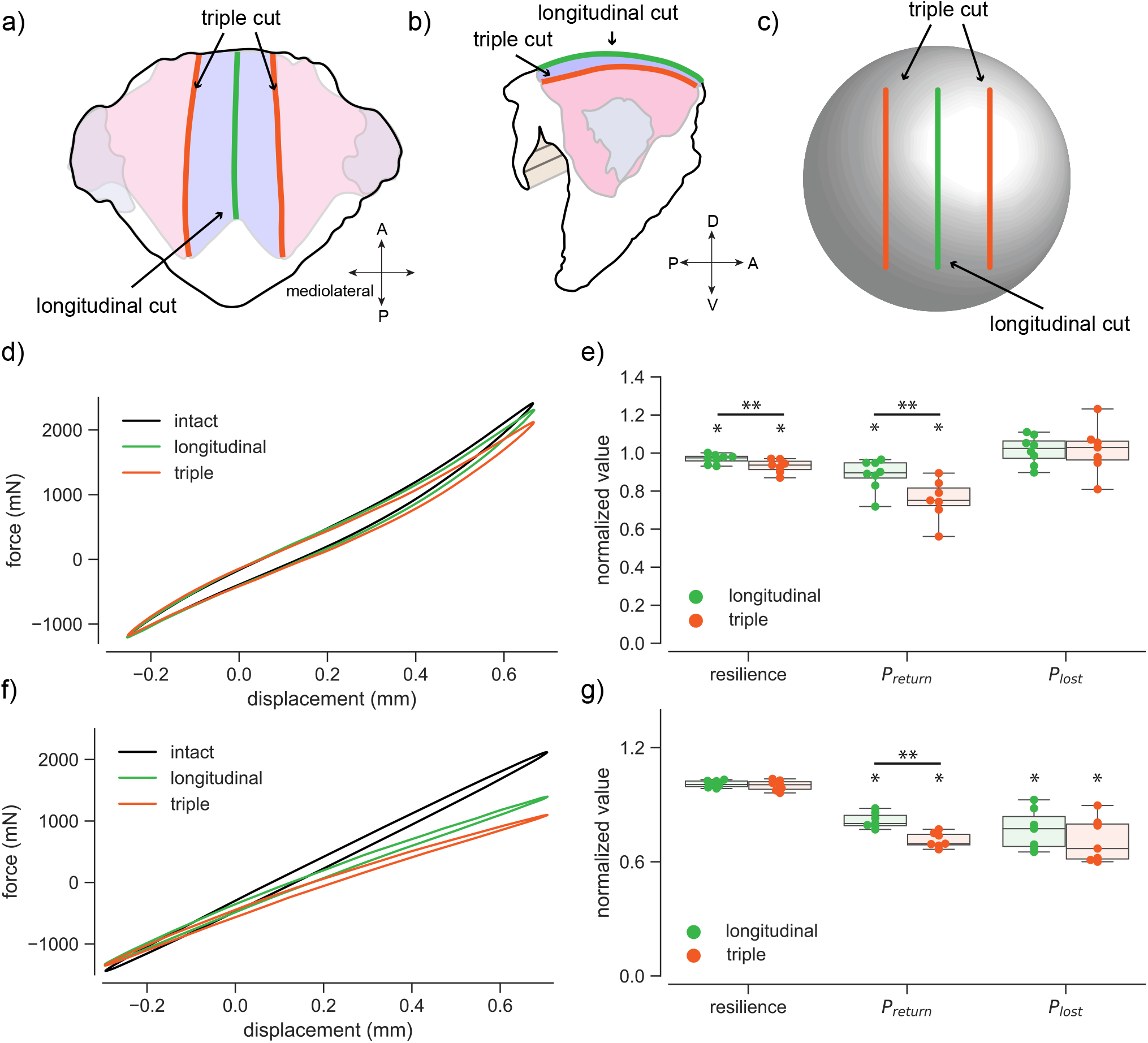
Strain disruption experiments. Dorsal a) and lateral b) views of the second thoracic segment. In addition to the features described in Fig. 6, the green line marks the longitudinal cut condition while red lines mark the triple cut. c) analogous dorsal view of a ping pong ball highlighting the cuts labeled in a). d) Average force-displacement plots for an intact, longitudinal cut, and triple cut thorax. At each time point, we plot the average force against the average displacement for each condition. Positive displacement denotes compression. e) Effects of strain disruption on resilience, *P_return_*, and *P_lost_* in thoraxes with passive musculature and a longitudinal cut (N = 8) or triple cut (N = 7). All data was normalized to the paired, intact thorax and taken at an oscillation frequency of 25 Hz. Boxplots denote the mean, quartiles, and range of the raw data while circles are individual data points. Single * denote a significant difference from the paired, intact thorax while ** mark differences between conditions. f) Same as d) but for ping pong balls. g) Same measures as e) but for ping pong balls with longitudinal cuts (N = 7) or triple cuts (N = 7).

For the ping pong ball, the isolated scutum (N = 5) had a *P_return_* of only 25.6 ± 6.5% (*p* = 0.002) of the intact system and a and *P_lost_* of 28.5 ± 3.5% (*p* < 0.001). Similarly, the isolated joint (N = 7) had a *P_return_* of 27.7 ± 5.1% (*p* = 0.001) and *P_lost_* of 22.8 ± 2.5% (*p* = 0.001). Notably, strain coupling between ping pong ball regions affects both *P_return_* and *P_lost_* because the sum of the joint and scutum regions is less than 1 for both measures.

The near monotonic decrease in *P_return_* and *P_lost_* led to normalized resilience values of 99.0 ± 2.6% for the isolated scutum and 102.7 ± 2.0% for the isolated joint. There was no statistically significant difference between the isolated scutum and the intact ping pong ball (*p* = 0.474), but resilience for the isolated joint significantly increased (*p* = 0.015). Although there is a slight increase in resilience in the ping pong ball under the isolated joint condition, these results indicate that the mechanical properties of both regions are nearly identical to that an intact ping pong ball.

Unlike the intuition developed from the ping pong balls, isolated thorax regions see sharp decreases in spring performance. The isolated scutum (N = 8) and isolated joint (N = 7) had normalized resilience values of only 76.6 ± 4.1% (*p* < 0.001) and 82.5 ± 3.6% (*p* < 0.001), respectively. The reduction in resilience is evident in the non-proportional changes in *P_return_* and *P_lost_*. Similar to the ping pong ball, when we isolated the scutum, *P_return_* fell to 21.9 ± 5.5% (*p* < 0.001) but *P_lost_* only decreased to 56.3 ± 14.4% (*p* < 0.001). When we isolated the joint, *P_return_* for the isolated joint was 47.5 ± 9.1% (*p* < 0.001) of the intact thorax. Contrary to expectations, *P_lost_* remained unchanged (*p* = 0.349). This suggests that the majority of energy dissipation occurs in the wing joints and the addition of a scutum therefore provides free *P_return_*. Similar to the ping pong balls, *P_return_* benefits from mechanical coupling between regions.

### 3.4 Disruption of thoracic shape reduces energy exchange performance

Isolating thorax regions alters bulk mechanics, but is is unclear whether these effects are due to changes in strain propagation or removal of material. Therefore, we performed a series of manipulations that disrupt strain but do not remove material. In the thorax, the longitudinal cut disrupts the transverse arch, while the triple cut disrupts strain between the wing joint and scutum.

For ping pong balls, disrupting the transverse arch via a longitudinal cut (N = 7) led to a *P_return_* of 81.7 ± 3.8% of the intact system (*p* = 0.001) and *P_lost_* of 74.2 ± 9.8 % (*p* = 0.013). When we added two parallel cuts for the triple cut condition (N = 7), *P_return_* was only 71.9 ± 3.6% (*p* < 0.001) of the intact ping pong ball and *P_lost_* dropped to 68.1 ± 10.8 % (*p* = 0.012). Similar to the region removal experiments, the nearly proportional decrease in *P_return_* and *P_lost_* led to normalized resilience values identical to the intact ping pong ball for both the longitudinal (*p* = 0.243) and triple (*p* = 0.935) cut conditions. This indicates that changes in strain propagation does not affect spring performance.

Similar to the ping pong balls, altering strain without removing material reduced *P_return_* in thoraxes. However, *P_lost_* was unchanged. For thoraxes, the longitudinal cut (N = 8) reduced *P_return_* to 88.1 ± 7.6% (*p* = 0.010) while *P_lost_* was unaffected (*p* = 0.803). Similarly, the triple cut (N = 7) reduced *P_return_* to 75.1 ± 9.9% (*p* = 0.002) while *P_lost_* remained constant (*p* = 0.984). Unlike the ping pong ball, changes in strain propagation did not affect *P_lost_*. A constant *P_lost_* but reduced *P_return_* led to normalized resilience values of 96.9 ± 2.3% (*p* = 0.008) for the longitudinal cut and 93.1 ± 3.4% (*p* = 0.002) for the triple cut condition. These results indicate that strain propagation is critical for thorax mechanics because strain disruption without the removal of material significantly alters bulk mechanical properties.

## 4 Discussion

### 4.1 Frequency dependent consequences of frequency independent thorax mechanics

The thorax’s frequency response is an important mechanical property for flight because insects may utilize wingbeat frequency modulation to overcome flapping wing instabilities [27, 28]. In air, oscillating systems typically exhibit frequency independent damping [24]. However, the introduction of fluids causes velocity dependencies. For instance, isolated, passive muscle exhibits strong frequency dependent damping, in part due to its high water content [29, 30, 31]. However, despite the presence of viscoelastic muscle, a cockroach leg is well characterized by a structural damping model, not a viscoelastic one [26]. Despite larger amounts of muscle, we find that the hawkmoth’s exoskeleton still dominates thorax properties, behaves as a structurally damped material, and leads to frequency independent mechanics (Fig. 4 b, c, & d) like in the cockroach leg.

The interactions of a frequency independent thorax with frequency dependent power requirements bounds the range of energetically preferable wingbeat frequencies. Unlike *P_return_* and *P_lost_*, both *P_inertial_* and *P_aero_* scale cubically with wingbeat frequency [5]. Because *P_return_* is frequency independent, there is only one wingbeat frequency that minimizes *P_inertial_* via elastic energy exchange. At higher wingbeat frequencies, the energy reduction from elastic energy exchange becomes negligible. Although moths do not possess a clutch mechanism in the wing joint, it is unclear whether this energetically optimal frequency corresponds to a true resonance because of nonlinear aerodynamic damping, lack of accurate estimates of effective wing mass, and uncertainties about transmission ratio between the muscles and wings. Unlike a viscously damped system, *P_lost_* places a lower bound on wingbeat frequencies. In this regime, a larger portion of energy expenditure would be spent overcoming thorax dissipation.

Beyond just hawkmoths, frequency independent damping may be ubiquitous in insect joints. At many length scales, components of the insect flight apparatus have frequency independent mechanical properties. Proteins [17], composite structures [32], and now our data on *Manduca sexta* thoraxes all demonstrate frequency independent damping. Extending beyond insect flight, both resilin and chitin are found in the spring-like structures of many arthropods [33]. Although Burrows *et al.* did not perform frequency sweep experiments, they found that froghopper insects store elastic energy in a composite structure composed predominantly of chitin and resilin [34]. In addition, resilin is found in the joints of cockroach legs [35], which is known to exhibit frequency independent damping [26]. The building blocks of insect joints are structurally damped, which is maintained as the systems are built up into an entire flight apparatus. Perhaps frequency independent damping is a ubiquitous property of insect joints.

While the structural damping model accurately represents harmonic thoracic deformations, the model is non-physical. Under harmonic oscillations, the imaginary *γ* term manifests as a phase lag between force and displacement. However, for non-harmonic conditions, the complex stiffness indicates that a non-physical complex force is required for a real displacement. In addition, the model is non-causal because it requires future states to predict its current state [36]. Therefore, the structural damping model cannot be used as an equation of motion to model oscillatory dynamics beyond harmonic oscillations, even if it provides a good phenomological fit for the behavior of the system. Mechanistic equations of motion for structurally damped materials (strain-dependent dissipation) remain a challenging frontier. Physically, frequency independent damping can occur if a system has many stress relaxation time constants [37]. At each oscillation frequency, the nearest time constant dominates, effectively leading to frequency independent damping. However, to the best of our knowledge these conditions have not yet been translated into a widely accepted mechanistic and mathematical formulation. Despite these limitations, a structural damping model provides a mathematical basis for investigating energy exchange under harmonic conditions.

### 4.2 Thoracic exoskeleton returns the maximum amount of beneficial elastic energy

The general approach to quantify elastic energy exchange is to estimate *P_total_* assuming first with no elastic energy exchange and then with perfect elastic energy exchange. The difference between the two *P_total_* values represents the maximum energetic benefit of elastic energy exchange. Using this general strategy in *Manduca sexta* but with different experimental approaches, Willmott & Ellington (blade element models), Sun & Du (computational fluid dynamics), and Warfvinge et al. (particle image velocimetry) concluded that elastic energy exchange can reduce *P_total_* by up to 20 - 35% [8, 7, 9]. To extend upon this work, Dickinson & Lighton utilized respirometric measurements combined with model based estimates of *P_aero_* and *P_inertial_* [5]. They found that 11% elastic energy exchange accounted for the differences in blade element model and respirometric estimates of *P_total_* in *Drosophila hydei*. However, these estimates rest on the assumption that muscle efficiency is constant. A following study shows muscle efficiency in the closely related *Drosophila melanogaster* can vary by three-fold depending on motor output [38]. In addition to modeling, Weis-Fogh reported preliminary measurements of the static relationship between wing angle and torque in locusts (*Schistocera gregaria*), privet hawk moths *Sphinx ligustri*, and dragonflies *Aeshna grandis* [39]. He concluded that the exoskeleton was stiff but was unable to quantify elastic energy exchange. In addition, it is unlikely that artificially backdriving the wings accurately reproduces thorax deformations seen *in vivo* because muscles are necessary to properly engage the wings into flight position [40].

We have found that the *Manduca sexta* thoracic exoskeleton returns 8.6 ± 2.4 W kg^−1^. Compared to computational fluid dynamics [41], robophysical [42], and kinematic estimates [8, 43] of body mass specific inertial power requirements 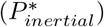, the passive thorax has the capacity to reduce 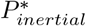 by 25 - 60% (Fig. 8). Our results agree with the previously discussed estimates of the maximum useful elastic energy exchange in *Manduca sexta*. For example, Willmott & Ellington estimated a maximum 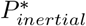 reduction 6 ± 3 W kg^−1^. In addition, Sun & Du predict that elastic structures return 9 W kg^−1^ [7]. Because the thorax itself returns the maximum beneficial energy, energy exchange from other structures or increased thorax stiffness would not affect flight power requirements. Therefore, exoskeletal deformations alone are sufficient for reducing *P_total_* to a minimum.

**Figure 8:**
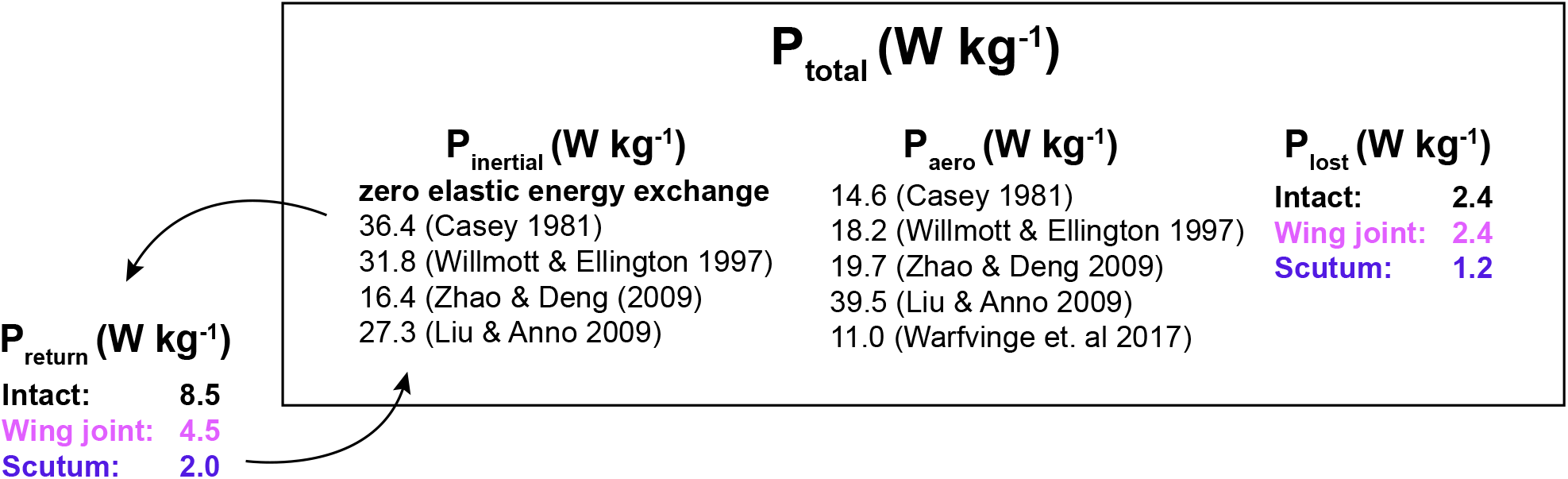
Power measurements during *Manduca sexta* flight. The total mechanical power produced by the muscles (*P_total_*) must supply *P_inertial_* and *P_aero_*. In addition, our results show that *P_lost_* is non-negligible and substantial recycling of *P_inertial_* via elastic energy exchange (*P_return_*).

Adding a stiff exoskeleton has potential detriments to insect flight. Although the thorax returns significant energy, it also dissipates 2.4 ± 0.5 W kg^−1^ that cannot be recovered (Fig. 8). In *Manduca sexta*, thorax elasticity leads to a net reduction in *P_total_*, but it is possible to dissipate more energy than recovered. In addition, before the wings have built up sufficient kinetic energy to deform the exoskeleton, insects must expend additional power to deform the thorax. Similarly, to rapidly accelerate the wings from a stop, *Manduca sexta* must now deform the stiff exoskeleton in addition to providing *P_inertial_* and *P_aero_*. To bring the wings to a sudden stop, *Manduca sexta* must rapidly dissipate stored elastic energy. However, for *Manduca sexta*, the benefits of minimizing *P_total_* during sustained, steady flight may outweigh these drawbacks.

These results provide an updated perspective on the power requirements of *Manduca sexta* flight. Due to experimental challenges in estimating *P_inertial_* and *P_aero_*, there is wide variability in the literature (Fig. 8) [43, 8, 42, 41, 9]. This variation has led some authors to argue that there is no benefit of elastic energy exchange because they find that *P_aero_* exceeds *P_inertial_* and the wing therefore has no excess kinetic energy [41, 42]. However, this argument depends on the time course of *P_inertial_* and *P_aero_* and not the wingbeat averaged values. In support of the elastic energy exchange hypothesis, our results show that the passive flight system has substantial energy exchange capacity. If the energy to deform the thorax does not come from excess wing kinetic energy, then *Manduca sexta* would need to provide this energy from muscles, which goes against an energy minimization strategy [44]. In addition, compared to the most recent and only direct measurement of power transferred to the surrounding air, dissipation in the thorax accounts for nearly 20% of power dissipation [9]. By dissipating substantial energy, losses in the thorax aid in wing deceleration, but reduce the maximum useful *P_return_*. Therefore, under the perfect energy exchange condition, *P_total_* must supply *P_lost_* in addition to *P_aero_*.

### 4.3 Strain coupling improves energy exchange performance

Elastic energy exchange is a common theme in biology, yet animals do not possess ideal springs [44, 2]. Instead, spring-like behavior arises from physical interactions across many length scales. For instance, the composite structure of invertebrate cuticle contributes to its material properties and some authors point to material testing results as a predictor of function [45, 17, 46]. However, a structure’s macroscopic shape can significantly alter bulk mechanics by coupling stretching and bending modes [47, 18]. For invertebrates, spring-like structures are frequently materially heterogeneous and possess complex geometry, which makes it difficult to determine the factors governing spring-like behavior [48, 49, 50]. Our dynamic mechanical testing experiments have identified a simple mechanical model of thorax mechanics, but it is unclear what properties of the multifunctional thorax give rise to its bulk mechanical properties.

To unravel the determinants of spring-like properties in the thoracic exoskeleton, we began by isolating two of the main functional thoracic regions and grounding our interpretation by comparing to a homogeneous hemisphere. We found that both ping pong balls and thoraxes exhibit shape dependent coupling of strain between spatially separated regions, which is evident in the nonlinear summation of *P_return_* between regions (Fig. 6 e & g). This strain coupling is also evident in thorax strain visualization (Fig. 2 d). As the phragma compresses the thorax (Fig. 1 b), the wing joint and scutum separate (Fig. 2 d). This separation leads to tension that couples the two regions. Unlike a homogeneous hemisphere, both isolated thorax regions had lower resilience than the intact thorax. This suggests that local mechanical properties depend on global structure, perhaps due to the thorax’s complex shape or material heterogeneity. To disentangle the effects of shape and material, we disrupted strain in both systems without removing material. We again observed that, unlike the homogeneous hemisphere, local resilience and therefore mechanical properties are dependent on global strain propagation (Fig. 7 e & g). In addition to expected changes in stiffness, we observe that altering strain patterns in the homogeneous hemisphere can affect dissipation, but does not in the thorax.

Although the exoskeleton is a single, continuous structure, functional specialization of thorax regions decouples *P_return_* and *P_lost_*. For instance, the wing joint must withstand the high strains necessary for wing movement whereas the scutum may be optimized for spring-like behavior [19]. Because dissipation occurs primarily in the wing joint, the addition of scutum deformations via indirect actuation increases *P_return_* by 50% but does does not increase *P_lost_*. The exoskeletal’s ability to return the maximum useful elastic energy may explain the general lack of tendons in insects with indirect musculature [45]. Although active muscle may recycle substantial energy, our results in tandem with previous estimates of the maximum useful elastic energy exchange suggest that energy exchange from other sources would not affect *P_total_* [8, 7, 9].

These results indicate that coupling between functional regions can have significant energetic consequences for biological systems. For instance, *Manduca sexta* benefits from the nonlinear summation of *P_return_*. Beyond insect flight, elastic energy exchange in exoskeletal structures is a key component of power amplified behavior [48, 49, 50, 2]. Like other biological structures with complex geometry, the performance of spring-like structures in the thorax may be highly dependent on their integration with the global system and cannot be characterized in isolation [51, 18].

## 5 Conclusion

For *Manduca sexta* and insects with indirect actuation, the power requirements directly related to wing movement (*P_inertial_* and *P_total_*) as well as those associated with thorax deformations (*P_return_* and *P_lost_*) are critical for flight energetics. At a minimum, an insect requires its wing joints to fly. By adding scutum deformations via indirect actuation, we find that *Manduca sexta* substantially increases elastic energy recovery. In many systems, such as a homogeneous hemisphere, increasing elastic energy recovery requires a proportional increase in energy dissipation. However, we find that elastic energy recovery and dissipation are decoupled in the *Manduca sexta* thorax, likely due to regional specialization of the exoskeleton. Because of this decoupling, there is no energetic cost to increasing *P_return_*. By adopting an indirect actuation strategy, insects can independently tune *P_return_* to minimize *P_total_*. Via dynamic mechanical testing, we also establish that the thorax’s mechanical properties are well approximated as a structurally damped, elastic material, which provides a useful characterization for integrated models of flapping wing flight in insects.

## 5.1 Acknowledgements

The authors thank James Lynch for helpful discussions.

## 5.2 Funding sources

This work was supported by U.S. National Science Foundation CAREER grant (Physics of Living Systems 1554790) to S.S. and the NSF Physics of Living Systems SAVI student research network (GT node grant — 1205878)

